# Neural basis of learning guided by sensory confidence and reward value

**DOI:** 10.1101/411413

**Authors:** Armin Lak, Michael Okun, Morgane Moss, Harsha Gurnani, Karolina Farrell, Miles J Wells, Charu Bai Reddy, Adam Kepecs, Kenneth D Harris, Matteo Carandini

**Affiliations:** Institute of Ophthalmology, University College London, London WC1E 6BT, UK; Institute of Neurology, University College London, London WC1E 6BT, UK; Centre for Systems Neuroscience, University of Leicester, Leicester LE1 7RH, UK; Cold Spring Harbor Laboratory, 1 Bungtown Road, NY 11724, USA

## Abstract

Making efficient decisions requires combining present sensory evidence with previous reward values, and learning from the resulting outcome. To establish the underlying neural processes, we trained mice in a task that probed such decisions. Mouse choices conformed to a reinforcement learning model that estimates predicted value (reward value times sensory confidence) and prediction error (outcome minus predicted value). Predicted value was encoded in the pre-outcome activity of prelimbic frontal neurons and midbrain dopamine neurons. Prediction error was encoded in the post-outcome activity of dopamine neurons, which reflected not only reward value but also sensory confidence. Manipulations of these signals spared ongoing choices but profoundly affected subsequent learning. Learning depended on the pre-outcome activity of prelimbic neurons, but not dopamine neurons. Learning also depended on the post-outcome activity of dopamine neurons, but not prelimbic neurons. These results reveal the distinct roles of frontal and dopamine neurons in learning under uncertainty.

## Introduction

Making efficient decisions requires combining present sensory evidence with previous reward values, and learning from the resulting outcome. It is not known, however, how the brain performs these computations. On the one hand, studies of perceptual decisions have established that observers carry estimates of sensory confidence, i.e. the probability that a percept is correct, and use it to estimate upcoming rewards (Gold and Shadlen, 2007; Kepecs et al., 2008; Kiani and Shadlen, 2009). On the other hand, studies of reward learning have shown how decisions are informed by past rewards, and formalized this process through reinforcement learning models based on predicted value and prediction error (Daw and Doya, 2006; Lee et al., 2012; Samejima et al., 2005; Schultz, 2015; Sutton and Barto, 1998). Animals and humans must be able to combine these computations, because their choices reflect both past rewards and present sensory stimuli (Fan et al., 2018; Feng et al., 2009; Hirokawa et al., 2017; Whiteley and Sahani, 2008). However, it is not known how neuronal signals combine sensory confidence and past rewards to predict upcoming rewards, and how these signals guide learning and inform subsequent choices.

A large body of evidence has shown that neurons in medial frontal cortex encode predicted value as inferred from past outcomes (Moorman and Aston-Jones, 2015; Otis et al., 2017; Pinto and Dan, 2015; Pratt and Mizumori, 2001; Rushworth et al., 2011). However, when decisions depend also on sensory evidence, predicted value should reflect not only the size of the reward, but also the confidence in the accuracy of the percept (Dayan and Daw, 2008; Lak et al., 2017). Predicted value should therefore be the product of these two factors. It is not known whether responses in medial frontal cortex reflect this computation, and thus appropriately encode predicted value.

A promising candidate for signals combining sensory confidence and past rewards is the prelimbic region of medial frontal cortex (PL). PL sends and receives projections from midbrain dopamine neurons (Beier et al., 2015; Carr and Sesack, 2000; Morales and Margolis, 2017). Lesion or inactivation of PL renders animals insensitive to reward value (Corbit and Balleine, 2003; Killcross and Coutureau, 2003; Ostlund and Balleine, 2005) and might impair sensory detection (Le Merre et al., 2018). In simple conditioning tasks, PL neurons may signal future rewards (Moorman and Aston-Jones, 2015; Otis et al., 2017; Pratt and Mizumori, 2001). It is thus possible that PL neurons compute the appropriate combination of sensory confidence and past rewards to predict upcoming rewards.

Another candidate for signals combining sensory confidence and past rewards are the dopaminergic neurons in midbrain. These neurons can encode predicted value and reward prediction error (Bayer and Glimcher, 2005; Cohen et al., 2012; Schultz et al., 1997). Their responses may also reflect sensory confidence (Lak et al., 2017). However, it is not known whether their activity appropriately combines reward values and sensory confidence. Moreover, while the causal roles of dopamine reward prediction errors in learning from past rewards is well established (Hamid et al., 2016; Kim et al., 2012; Parker et al., 2016; Stauffer et al., 2016; Steinberg et al., 2013; Tsai et al., 2009), it remains unknown whether dopamine responses account for the effect of decision confidence on learning.

Furthermore, the functional role of neuronal signals encoding predicted value remains untested. Theoretical models propose that predicted value is computed so that it can be compared to outcome and drive learning (Sutton and Barto, 1998). As such, the predicted value of the chosen option is not necessary for the decision itself (it is computed after the decision), but rather is necessary for learning. To test these possibilities, one should manipulate predicted value signals before decision outcome.

To address these questions, we developed a decision task that requires mice to combine past rewards with present sensory evidence. We devised a simple behavioral model that describes their choices, and makes strong predictions about how reward value and sensory confidence would affect the decision in the next trial. The model makes trial-by-trial estimates of predicted value and prediction error, both of which depend on confidence and past rewards. We found neural correlates of predicted value in the pre-outcome activity of PL neurons and of dopamine neurons of ventral tegmental area (VTA). We found neural correlates of prediction error in the activity of dopamine neurons at the time of outcome. Manipulating pre-outcome activity of PL neurons, but not dopamine neurons, drove learning. Conversely, perturbing the post-outcome activity of dopamine neurons, but not PL neurons, guided learning. These results reveal how frontal and dopaminergic circuits guide learning under sensory and value uncertainty.

## Results

We begin by describing the behavioral task and the model that fits the observed choices. We then establish correlates for the model’s internal variables in PL populations and in VTA dopamine neurons, and demonstrate their specific, causal roles in learning.

### Behavioral signatures of learning guided by sensory confidence and reward value

To study decisions guided by sensory signals and reward values, we developed a task for head-fixed mice (Figure 1A-C). We presented a grating of variable contrast on the left or right side and the mouse indicated the grating’s position by steering a wheel with its forepaws (Figure 1A) receiving water for correct responses (Figure 1B) or a noise sound for incorrect ones (Burgess et al., 2017). We manipulated the reward delivered for correct left and right choices, with one side receiving twice as much water (2.4 vs. 1.2 μl); the more-rewarded side switched without warning in blocks of 50-350 trials and was not otherwise cued (Figure 1C).

**Figure 1.**
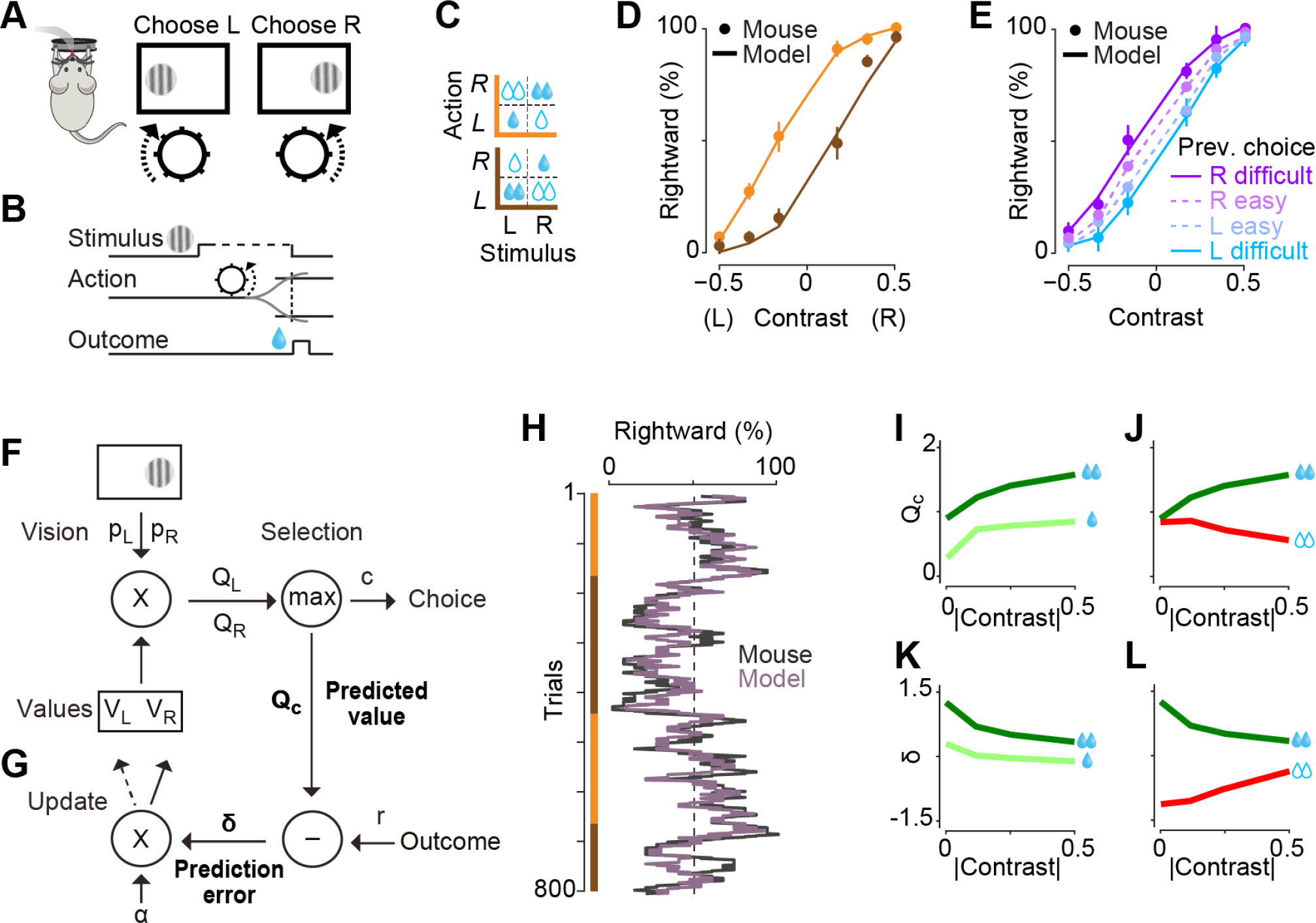
Behavioral and computational signatures of decisions guided by reward value and sensory confidence. **A,B**)Schematic of the 2-alternative visual task. **C**) Rewards for correct choices were higher on the right side (orange) or on the left side (brown), with the more-rewarded side switching in blocks of 50-350 trials. **D**) Behavior of an example mouse in blocks with large reward on right (orange) or left (brown). Curves are predictions of the behavioral model in (F-G). Error bars show standard error of the mean (s.e.) across trials. See Figure S1B-D for similar results from all mice, and for reaction times. **E**) Behavior of the same mouse as a function of the difficulty of previous rewarded trial, for difficult choice (low contrast) or easy choice (high contrast) to the left or right. Curves are predictions of the model presented in (F-G). See Figure S1E for all mice, and Figure S1F,G for similar results in a task without reward manipulation. **F,G**) Behavioral model of choice (F) and learning (G). **H**) Running average of probability of choosing right, in a session containing four blocks (orange vs. brown). Black: mouse behavior. Light purple: model predictions. **I**) Averaged estimates of *Q*_*C*_ as a function of absolute contrast (i.e. regardless of side), for correct decisions towards the large-reward side (dark green) and correct decisions towards the small-reward side (light green). **J**) Averaged estimates of *Q*_*C*_ for correct decisions (dark green) vs. incorrect decisions (red), both made towards the large-reward side. See Figure S1K for errors towards small-reward side. **K-L**) Similar to (I-J) but for reward prediction error δ.

Mice performed this task proficiently, integrating past rewards and current sensory evidence when making decisions (Figure 1D). In this task, efficient decisions depend on a combination of sensory uncertainty and reward value. High contrast stimuli are unambiguous, and should always be chosen, because choosing the other side would give no reward. Conversely, in low contrast trials, decisions should favor the side paired with larger reward, as can be derived mathematically (Whiteley and Sahani, 2008) and with simulations (Figure S1A). Mice mastered the task, efficiently combining sensory evidence and past rewards (Figure 1D; Figure S1B for n=10 mice). Their psychometric curves shifted sideways between blocks (Figure 1D, Figure S1B), so that reward value predominantly affected decisions for low-contrast stimuli (*P* < 10^−10^, 1-way ANOVA).

Despite this good performance, however, the mice were gradual, and hence suboptimal, in their learning of reward value. Because we switched the side associated with large reward across blocks of trials, it should be possible to infer the onset of a block from a single rewarded trial. Instead, the mice showed gradual learning following a block switch. For instance, they took ~12 trials to shift their low-contrast decisions by 10% towards the side newly paired with the larger reward (Figure S1C).

Mice were also suboptimal in another respect: their decisions depended on the decision confidence in the previous trial (Figure 1E). After a correct trial, the psychometric curve shifted towards the chosen side if that trial had been difficult (low contrast) but not if it had been easy (high contrast, Figure 1E). These results could not be explained by the presence or absence of rewards (only rewarded trials were included in the analysis), by a win-stay strategy, or by the block structure of the task (the analysis was performed within blocks). Similar results were seen across 8 mice (Figure S1E; Difficult: *P* = 0.01, Easy: *P* = 0.56, 1-way ANOVA). In fact, this effect of past decision confidence on future choices was also present in purely visual decisions, i.e. without manipulation of reward value (Figure S1F, G; Difficult: *P* = 0.01, Easy: *P* = 0.12, 1-way ANOVA).

Taken together, these results show that decisions in this task are jointly informed by available sensory evidence, by past reward values, and by past decision confidence. While the psychometric shifts due to reward size are beneficial for maximizing rewards, the effect of past stimulus on decisions is detrimental, because stimuli were presented in random order, so they had no bearing on the next trial.

### A model for decisions and learning under uncertainty

To describe mouse behavior and make testable predictions about its neural basis, we used a model that incorporates sensory uncertainty with conventional temporal difference reinforcement learning (Figure 1F,G). The model selects a choice and estimates its predicted value by combining sensory evidence with learned values (Figure 1F). In each trial, vision provides (noisy) probabilities *p*_*L*_ and *p*_*R*_ that the stimulus is on the left or right side. Multiplying these probabilities with learned values of left and right actions, *V*_*L*_ and *V*_*R*_, provides the expected values of left and right choices: *Q*_*L*_ = *p*_*L*_ *V*_*L*_ and *Q*_*R*_ = *p*_*R*_*V*_*R*_. The higher of these two determines the choice *C* (either *L* or *R*), its associated confidence *p*_*C*_, and its predicted value *Q*_*C*_ = *p*_*C*_ *V*_*C*_. Following the outcome, the model learns by updating the value of the chosen action by *V*_*C*_ ← *V*_*C*_ + *αδ*, where *α* is a learning rate, and *δ* = *r* − *Q*_*C*_ is the reward prediction error, i.e. the difference between available and predicted reward.

This behavioral model accounted quantitatively for the animals’ decisions (Figure 1D, E, H). It fitted the shift in psychometric curves due to reward size (Figure 1D, *curves*), predicted trial-by-trial decisions (Figure 1H, *purple trace*), and captured the time course of learning after block changes (Figure S1C). The model also accounted for the effect of past decision confidence on subsequent choices (Figure 1E, *curves*; Figure S1E-G, *curves*). Indeed, the prediction errors that drive learning in the model are larger when predicted value *Q*_*C*_ is smaller, as is the case at low contrast (where decision confidence *p*_*C*_ is low, because p_L_ ≈ p_R_ ≈ 0.5). Cross-validation confirmed the necessity of each model parameter; the full model (with no dropped parameters) provided the best fit in 8 out of 10 mice (Figure S1I, J).

Conversely, an alternative class of model that fully leverages the structure of the task (“model-based observer”) did not provide adequate fits (Figure S1M-Q). An observer that knows that only two reward sizes are available and that they occasionally switch side would only need to monitor whether a switch has occurred (Figure S1M). This observer, however, would not exhibit the dependence of choices on decision confidence in previous trials (Figure S1N-Q).

Having established the validity of our behavioral model in capturing the mice’s choices, we examined its testable predictions for two key internal variables, *Q*_*C*_ and *δ* (Figure 1I-L). For correct trials, the predicted value *Q*_*C*_ increases with stimulus contrast and is higher when stimuli appear on the large-reward side (Figure 1I). It is also higher for correct trials than for error trials, because in correct trials the decision confidence *p*_*C*_ tends to be higher (Figure 1J; Figure S1K). The reward prediction error *δ*, the difference between outcome and *Q*_*C*_, is larger following a larger reward, and decreases with stimulus contrast in correct trials (Figure 1K) but not in error trials, again reflecting the difference in decision confidence across these trials (Figure 1L, Figure S1L).

### Prelimbic neurons encode confidence-dependent predicted value

We next sought to identify neural correlates of the predicted value of choice *Q*_*C*_ in the activity of prelimbic region of medial frontal cortex (PL, Figure 2A). We used high-density silicon probes to record from 1,566 PL neurons in 6 mice. Of these, 316 neurons were significantly modulated by at least one task event (signed rank test on responses prior and after each task event, *P* < 0.01). A typical neuron showed a slight increase in firing rate following the stimulus and a large increase at the time of action, i.e. the onset of wheel movement (Figure 2B). Among the 316 task responsive neurons, most were modulated by action onset (78% of the neurons, *P* < 0.01, signed rank test), and smaller fractions by stimulus appearance (24%) or outcome delivery (19%, Figure 2C). Most neurons (54%) increased their firing prior to actions while others (24%) decreased their firing (Figure 2C; *P* < 0.01, signed rank test, n = 130-1080 trials depending on session).

**Figure 2.**
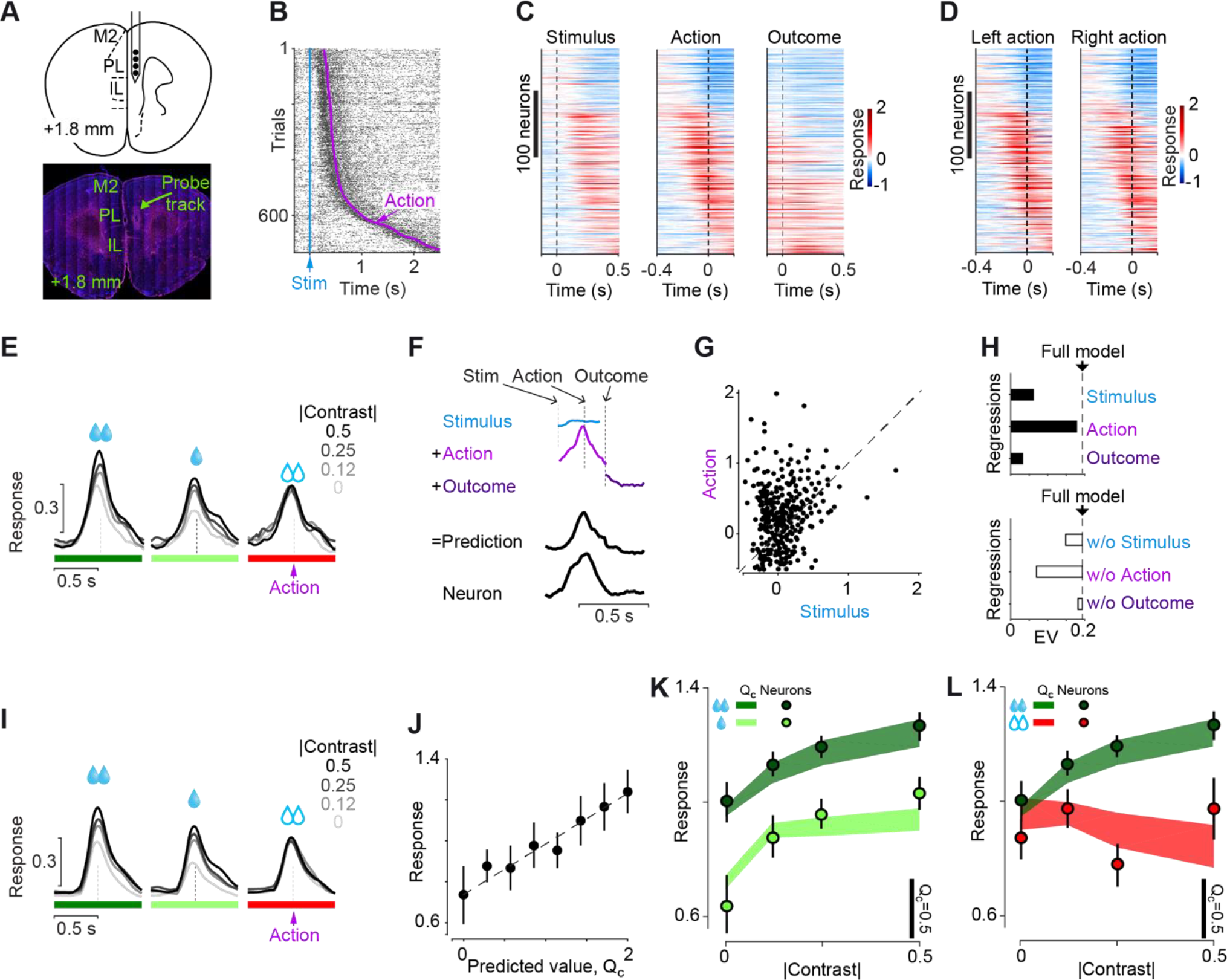
Prelimbic neurons encode confidence-dependent predicted value. **A**) Histological image showing the high-density silicon probe track in PL. **B**) Raster plot showing spikes of an example PL neuron, aligned to the stimulus onset (blue line) with trials sorted by action onset (purple dots). **C**) Responses of all task-responsive neurons (n=316), aligned to the time of stimulus, action, or outcome, sorted according to the time of maximum response in the middle panel. Responses were z-scored and averaged over all stimulus contrasts and possible outcomes. **D**) Same as the middle panel of (C), for trials with maximum stimulus contrast with Left or Right actions. **E**) Mean z-scored population activity triggered on action onset for correct choices towards the large-reward side (left), correct choices towards the small-reward side (middle), and incorrect choices towards the large-reward side (right). Responses for incorrect choices towards the small-reward side were smaller (P = 0.015, signed rank test, not shown) but such trials were rare. See Figure S2A for responses shown separately for neurons activated or suppressed at the time of the action. See Figure S2B for population activity triggered on outcome onset. **F**) The regression analysis estimates a temporal profile for each task event, which in each trial is aligned to the event onset time and scaled by a coefficient. The results are summed to produce predicted traces. **G**) The size of action and stimulus profiles for the full regression. Each dot presents one neuron. The responses are in units of standard deviation as responses of each neuron were z-scored. **H**) Top: cross-validated explained variance (EV) averaged across neurons for the full regression (dotted line), and for regressions each including only one type of event (bars). Bottom: variance explained by full regression (dotted line), and regressions each excluding one of the events (bars). **I**) Predictions of the regression only including action events triggered on action onset, as a function of stimulus contrast and trial type. **J**) Average action responses (estimated by regression on PL activity) as a function of trial-by-trial decision value *Q*_*C*_ (estimated from the behavioral model). **K**) Average action responses in correct trials as a function of stimulus contrast and reward size. Circles: mean; error bars: s.e. across neurons; shaded regions: model estimate of *Q*_*C*_. **L**) Same as (K) but for correct and error trials to the large-reward side. In J-L, only neurons with significant action profile were included (241/316 neurons). See Figure S2F,G for responses of remaining neurons.

Several aspects of PL activity around the time of the action were consistent with a signal encoding predicted value (Figure 2D,E). First, most PL neurons (95%) did not respond differently for the left and right actions (*P* < 0.01, signed rank test, Figure 2D). Second, PL activity depended both on stimulus contrast and on upcoming outcome (Figure 2E). In correct trials (Figure 2E, *green*), activity around action onset (−200 to 50 ms window) was higher when stimuli had higher contrast (*P* = 10^−6^, 1-way ANOVA) and were associated with larger rewards (*P* = 0.009, signed rank test). In error trials (red), which were most common in decisions towards the large-reward side, PL activity was lower than in correct trials (Figure 2E, *red P* = 10^−8^, signed rank test) and was not significantly modulated by contrast (*P* = 0.24, 1-way ANOVA). The effects of stimulus contrast, reward size, and correct/error were specific to neurons activated by action, and absent in neurons suppressed by action (Figure S2A).

The trial-by-trial activity of PL neurons quantitatively matched the predicted value of choice *Q*_*C*_ estimated by the behavioral model (Figure 2F-L). To show this, we used regression to estimate each neuron’s activity as a sum of temporal profiles corresponding to stimulus, action, and reward, with the magnitude of these profiles, but not the shape, allowed to vary between trials (Figure 2F). The profile for each event determines the average response of the neuron to that event, and the trial-by-trial magnitude estimates the neuron’s responses to the event on each trial. This analysis confirmed that in most neurons the response related to action was larger than responses related to stimuli and outcomes (Figure 2G, P = 0.0001, signed rank test). In most neurons (241/316) action events were sufficient to explain the observed neuronal activity (Figure 2H,I, Figure S2C-E). Here we focus on these neurons (see Figure S2F,G for the properties of the remaining neurons). Trial-by-trial variations in action-related activity correlated strongly with *Q*_*C*_ (Figure 2J, R^2^=0.88, *P* = 10^−4^, linear regression). Similar to *Q*_*C*_, PL activity increased with the size of the pending reward. Moreover, it increased with stimulus contrast (Figure 2K, *dark green* and *light green*), and it did so only for correct decisions (Figure 2L). Trial-by-trial variations in PL activity correlated better with *Q*_*C*_ than with measures of movement vigor such as wheel acceleration both at the population level and in individual neurons (Figure S2H; population: *P* = 0.001, signed rank test; 54 vs 22 neurons, *P* < 0.01, linear partial correlation). As we will see, optogenetic manipulations further support this observation. We conclude that *Q*_*C*_ is encoded in the pre-outcome activity of PL neurons.

### VTA dopamine neurons encode confidence-dependent predicted value and prediction error

To examine the integration of sensory evidence and reward value in the activity of VTA dopamine neurons, we measured their responses during the task using fiber photometry of GCaMP6 signals (Figure 3A,B). To allow sufficient time to measure Ca^2+^ transients, we modified the task slightly, and trained mice to respond after an auditory go cue that followed the visual stimulus (Figure 3B).

**Figure 3.**
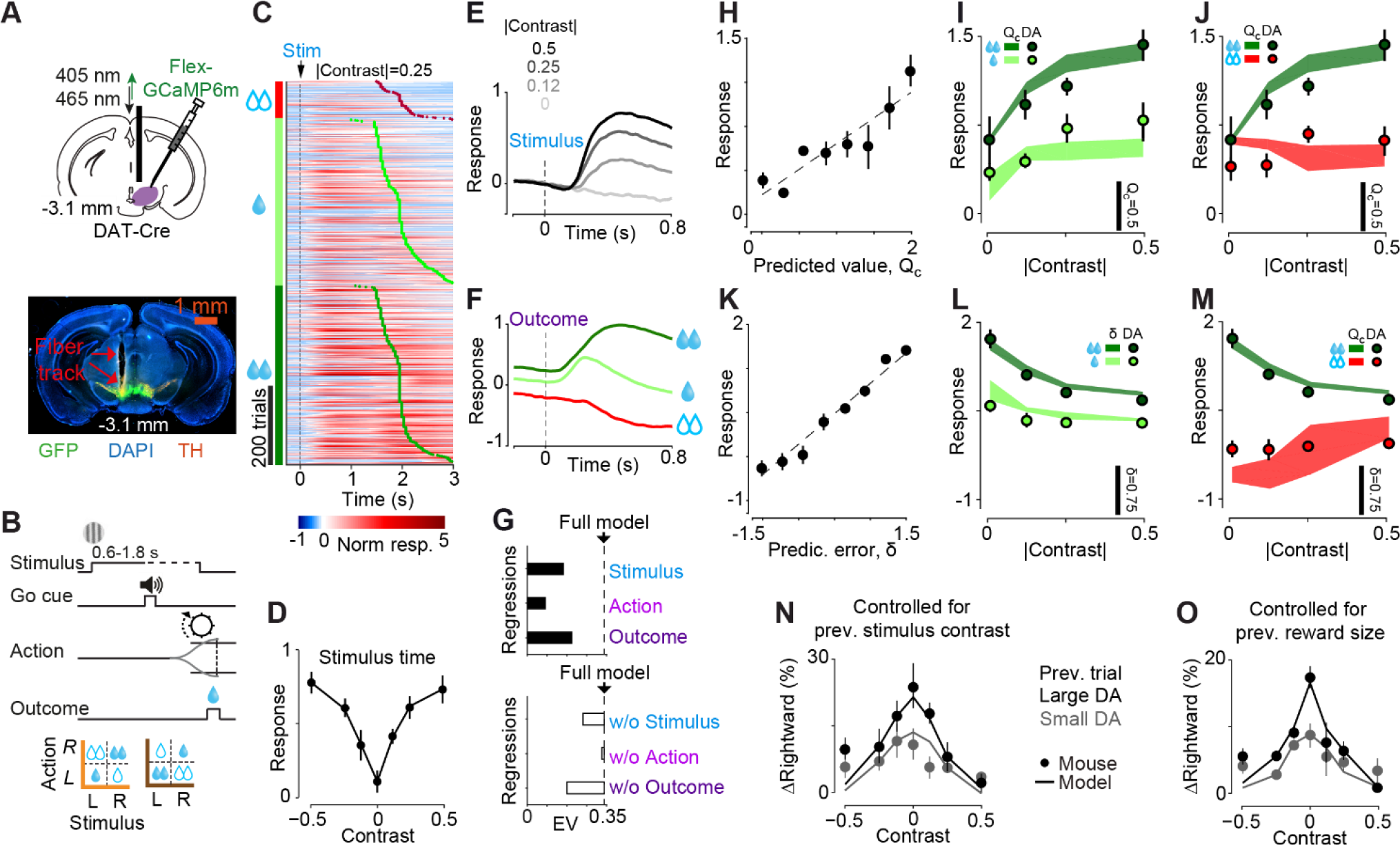
VTA dopamine neurons encode confidence-dependent predicted value and prediction error. **A**) Top: schematic of fiber photometry in VTA dopamine neurons. Bottom: example histology showing GCaMP expression and the position of implanted fiber above the VTA. **B**) Task timeline. To allow sufficient time for GCaMP measurement, decisions could be reported only after an auditory go cue. **C**) Trial-by-trial dopamine responses from all sessions of an example animal for trials with |contrast| = 0.25, aligned to stimulus onset (dashed line) and sorted by trial type (left column) and outcome time (red, light green and dark green dots). **D**) Average dopamine responses on correct trials as a function of contrast, for stimuli presented on the left or right side of the monitor. **E**) Population dopamine responses (n=5 mice) aligned to the stimulus. **F**) Population dopamine responses aligned to the outcome. **G**) Top: cross-validated explained variance (EV) averaged across mice for the full regression (dotted line), and for regressions each including only one type of event (bars). Bottom: EV of full regression (dotted line), and regressions each excluding one of the events (bars). **H**) Stimulus responses, estimated from regression, as a function of trial-by-trial decision value *Q*_*C*_, estimated by the behavioral model. **I**) Average stimulus responses in the correct trials as a function of stimulus contrast and trial type (error bars: s.e. across animals); shaded regions: model predictions of *Q*_*C*_. **J**) Same as (I) but for correct and error trials in which the large-reward side was chosen. **K**) Outcome responses, estimated from regression, as a function of trial-by-trial prediction error δ, estimated by the behavioral model. **L,M**) Same as (I, J), for outcome responses and model estimates of δ. **N**) Changes in the proportion of rightward choices (∆Rightward) as a function of dopamine activity to reward in the previous trial (black and gray: larger and smaller than 65 percentile, respectively), computed for each level of sensory stimulus in the previous trial (for left and right blocks separately), and then averaged. ∆Rightward reflects the difference between psychometric curve following rightward decisions and the psychometric curve following leftward decisions. **O**) Changes in the proportion of rightward choices as a function of dopamine activity to reward in the previous trial, computed for each reward size in the previous trial (for left and right blocks separately), and then averaged.

Dopamine activity was modulated both at the time of stimulus onset and at the time of outcome (Figure 3C-F). Following stimulus presentation, dopamine activity increased with the size of pending reward (Figure 3C, *P* < 0.004 in 5/5 mice, signed rank test) and with stimulus contrast (Figure 3D,E, *P* < 10^−4^ in 5/5 mice, 1-way ANOVA), regardless of stimulus side (Figure 3D, *P* > 0.08 in 5/5 mice, signed rank test). Dopamine activity was not significantly modulated at the time of actions (Figure S3, *P* > 0.13 in 5/5 mice, signed rank test). However, it was markedly increased at the time of outcome, especially after obtaining the larger reward (Figure 3C,F, *P* < 10^−4^ in 5/5 mice, 1-way ANOVA).

Both dopamine responses, following stimulus and following reward, closely matched the model variables estimated from behavior (Figure 3G-M). We used regression to estimate dopamine responses to stimulus presentation, action, and reward on every trial (Figure S3A). Omitting responses to action did not worsen the predictions (Figure 3G, Figure S3B,C), so we focused on responses to stimulus and outcome. At stimulus time, dopamine activity closely followed the behavioral model’s trial-by-trial estimates of predicted value *Q*_*C*_ (Figure 3H; population: R^2^=0.83, *P* = 0.001 and R^2^>0.57, *P* < 0.01 in 5/5 mice, linear regression), increasing with pending reward size and stimulus contrast for correct trials (Figure 3I), but not for incorrect trials (Figure 3J). At outcome time, moreover, dopamine activity closely followed the model’s estimates of prediction error *δ* (Figure 3K; population: R^2^=0.97, *P* = 10^−6^ and R^2^>0.88, *P* < 10^−4^ in 5/5 mice, linear regression). It increased with reward size and depended on the contrast of a stimulus that was no longer on the screen, decreasing with contrast in correct trials (Figure 3L), and not in error trials (Figure 3M). Thus, dopamine responses prior to outcome reflected *Q*_*C*_, in a manner similar to prelimbic activity, and dopamine responses after the outcome encoded *δ*.

The behavioral model also correctly predicted that dopamine responses at the time of outcome influenced subsequent choices (Figure 3N,O). If a choice (leftward or rightward) was followed by a large dopamine response, mice were more likely to make the same choice in the next trial. This effect could not be explained solely by the contrast of the previous stimulus (Figure 3N; *P* = 0.0002, 1-way ANOVA,) or by the size of the previous reward (Figure 3O; *P* = 0.0007, 1-way ANOVA,). The behavioral model captured it because its computation of the prediction error *δ* depends on both stimulus contrast and reward size.

### Pre-outcome activity of prelimbic neurons drives learning

Having established that PL signals prior to outcome closely correlate with estimate of predicted value, *Q*_*C*_, we asked whether these signals play a causal role (Figure 4). In our model, *Q*_*C*_ is determined only after making the choice *C*. The model thus predicts that reducing *Q*_*C*_ cannot influence the choice. Rather, reducing *Q*_*C*_ should affect learning, thus influencing subsequent trials. We tested these predictions through optogenetic inactivation, by directing brief laser pulses through an optical fiber in mice expressing Channelrhodopsin-2 (Chr2) in *Pvalb*-expressing inhibitory neurons of PL (Figure 4A,B, Figure S4A) (Guo et al., 2014; Olsen et al., 2012).

**Figure 4.**
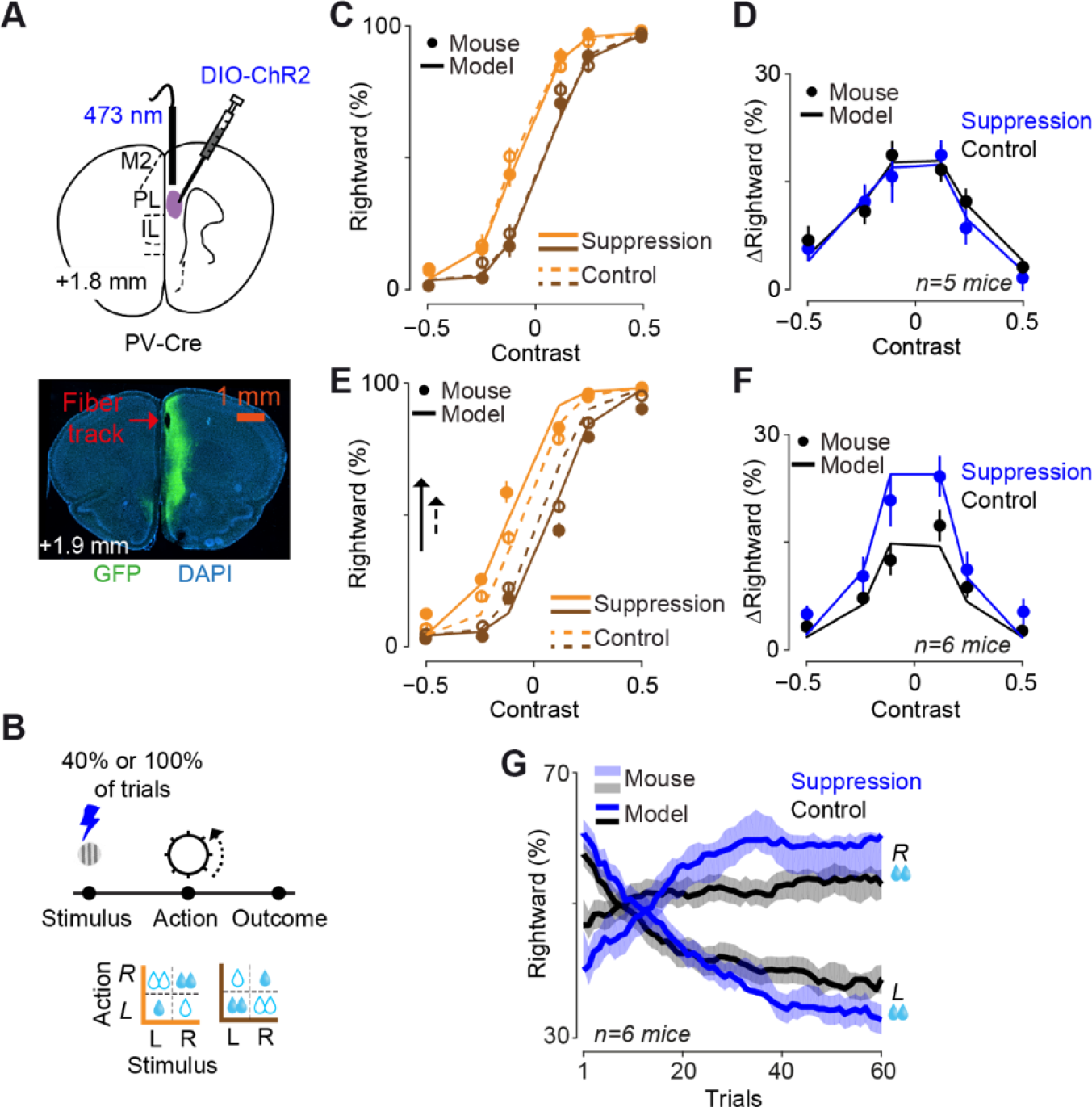
Learning, but not ongoing choice, depends on pre-outcome activity of prelimbic neurons. **A**) Top: to suppress PL population activity, we optogenetically activated Pvalb neurons. Bottom: example histology showing ChR2 expression in medial frontal cortex, and position of implanted fiber above PL. **B**) Inactivation occurred for 450 ms following stimulus onset in either 40% of randomly-selected trials (C,D) or in blocks of trials, forming four possible blocks (with or without suppression each with large reward on the left or right) (E-G). **C**) Reducing *Q*_*C*_ in the model does not influence ongoing choices. Curves are model predictions for trials with reduced *Q*_*C*_ (solid) and control trials (dashed). Consistent with the model prediction, suppressing PL neurons did not influence the performance in current trials. The data points show an example animal. See Figure 4SB for similar results in a task with no reward manipulation. **D**) Effect of PL suppression on psychometric shifts in 5 mice. Data points show the difference in the proportion of rightward choices between the L and R blocks of the control and suppression conditions (i.e. subtracting filled brown from filled orange circles, and subtracting empty brown from empty orange circles shown in (C)). Curves illustrates model fits on the data. **E**) Reducing *Q*_*C*_ in the model magnifies reward-induced psychometric bias. The arrow indicates the difference in the probability of rightward choice in trials with low contrast in the control (dashed) and in blocks with reduced *Q*_*C*_ (solid). Consistent with the model prediction, suppressing PL neurons during the task magnified the reward-induced side shifts of psychometric curves. The data points show an example animal. **F**) Effect of PL suppression on psychometric shifts in 6 mice. Curves illustrates model fits on the data (with reduced *Q*_*C*_ relative to control). **G**) The effect of PL suppression on trial-by-trial learning from the onset of block switch. The shaded areas indicate data (n=6 mice) in the control (black) and optogenetic suppression (blue) experiment, and curves are predictions of the model fitted on the data.

Consistent with the first prediction, suppressing PL did not influence ongoing decisions (Figure 4C,D). We suppressed PL activity in a subset of trials from the stimulus onset for 450 ms, and found no significant effect of suppression on the ongoing choices (*P* = 0.84, signed rank test). Similar results were observed in a simpler version of the task where reward sizes are equal and constant (Figure S4B).

Consistent with the second prediction, suppressing PL strikingly increased the effect of learning (Figure 4E-G). The model predicts that reducing *Q*_*C*_ in trials ending with reward would overestimate positive prediction error, *δ* = *R* − *Q*_*C*_, magnifying the subsequent shift in psychometric curves. We verified this prediction by suppressing PL activity from the stimulus onset in blocks of trials. As predicted, suppressing PL significantly increased the reward-induced shifts in psychometric curves (Figure 4E,F, *P* = 0.01, signed rank test). The model readily accounted for this effect (Figure 4E, *curves*) with the simple assumption that inactivation of PL subtracts a constant value from *Q*_*C*_ (Figure S4C). The model also closely predicted how inactivation of PL facilitated the progression of learning after the block switch (Figure 4G). These effects were not accompanied by changes in reaction time or in wheel acceleration (P = 0.43 and P=0.53, signed rank test). Also, these effects were seen only when suppressing PL activity before outcome. Consistent with the weak responses seen in PL at the time of outcome, suppressing PL at that time did not influence the choices (Figure S4D, *P* = 0.96, signed rank test). Taken together, these results indicate that PL causally encodes predicted value *Q*_*C*_. Pre-outcome activity of PL is necessary to learn from the upcoming outcome, and thus shape behavior in future trials.

### Learning relies on post-outcome, but not pre-outcome, activity of VTA dopamine neurons

Having observed that pre-outcome and post-outcome dopamine signals encode predicted value *Q*_*C*_ and prediction error *δ*, we next investigated their causal role (Figure 5). We implanted optical fibers above VTA in mice expressing Archaerhodopsin-3 (Arch3) or Chr2 in midbrain dopamine neurons (Figure 5A, Figure S5A) and we delivered brief laser pulses at the time of stimulus or of water reward.

**Figure 5.**
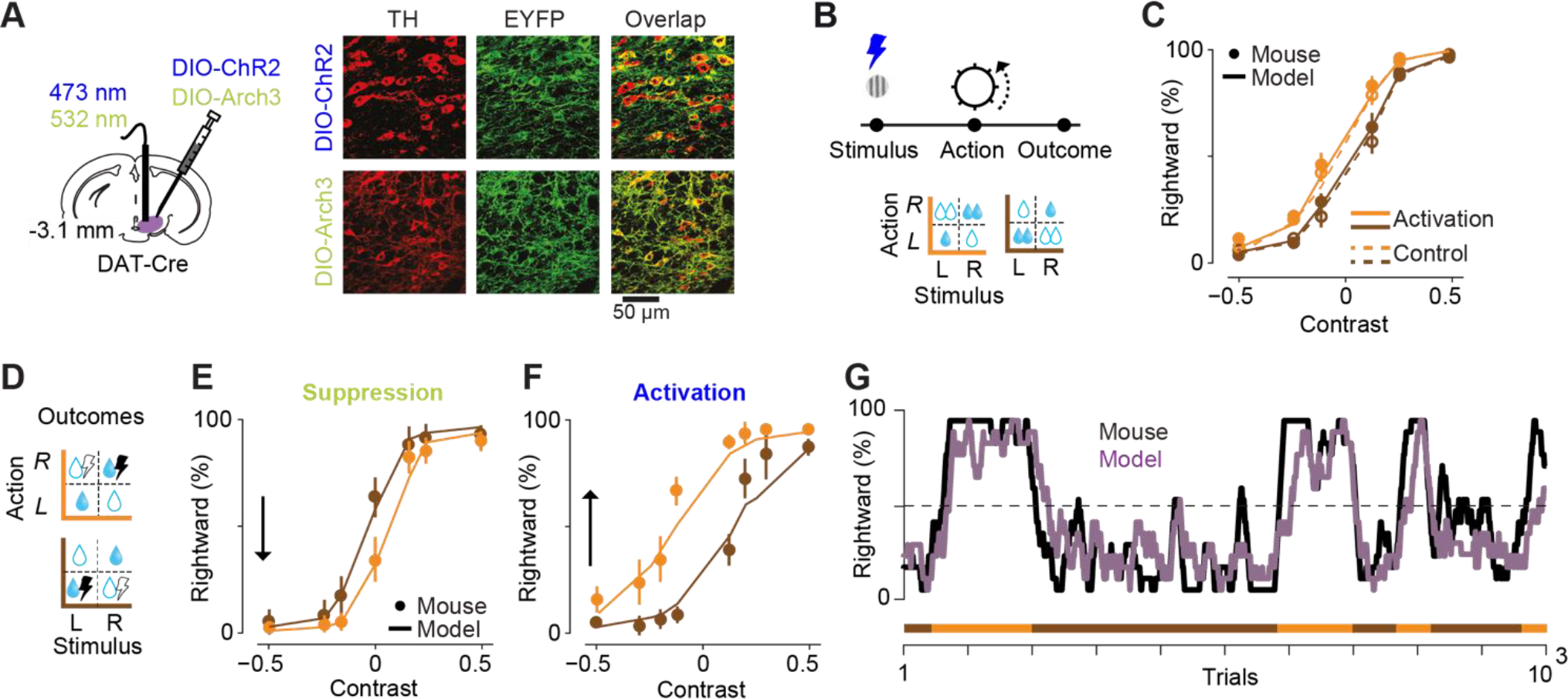
Learning relies on post-outcome, but not pre-outcome, activity of VTA dopamine neurons. **A**) Left: ChR2 or Arch3 were expressed in dopamine neurons and a fiber implanted over VTA. Right: Expression of ChR2 or Arch3 in dopamine neurons. **B**) In the first experiment light pulses were delivered at the time of visual stimulus in blocks of trials, forming four possible blocks (with or without activation each with large reward on the left or right). **C**) Behavior of an example animal in the activation trials (filled circles) and control trials (empty circles). Curves are model fits. Error bars are s.e. across trials. See Figure S5B for population data and Figure S5C-G for similar results in a task without reward manipulation or when activation started before the stimulus onset. **D**) Manipulation of dopamine responses at the time of outcome: light pulses were delivered following correct decisions towards one response side, which alternated in blocks of 50-350 trials. **E,F**) Model-predicted horizontal psychometric curve shift (curves) accounts for dopamine-induced behavioral changes (points). The arrow indicates the difference across blocks in the probability of rightward choice in trials with zero contrast. The psychometric shifts were independent of the hemisphere manipulated (P = 0.36, 2-way ANOVA). See Figure S5H-J for similar results across the population and reaction times. **G**) Running average of probability of rightward choice in an example session including 8 blocks (orange and brown). Black: mouse behavior. Purple: model prediction. See Figure S5K for averaged learning curves.

Manipulating dopamine activity prior to outcome influenced neither ongoing decisions nor learning in the subsequent trials (Figure 5B,C, Figure S5). Activating dopamine responses from the stimulus onset (for 450 ms) had no effect on the psychometric curves (Figure 5C; *P* = 0.46, signed rank test, Figure S5B). We observed similar results when we activated these neurons in a subset of trials in a simpler task without reward manipulation (Figure S5C-E). Also, activation starting prior to stimulus onset did not influence psychometric curves but slightly decreased reaction times (Figure S5F,G).

These results indicate that there is a fundamental difference between the signals encoding *Q*_*C*_ in the pre-outcome activity of PL and of VTA dopamine neurons. The former plays a causal role in learning but the latter does not.

In contrast, manipulation of post-outcome dopamine responses drove learning. We activated or suppressed dopamine responses during reward delivery for correct decisions towards one side, and alternated the side in consecutive blocks of trials (Figure 5D). The water rewards were equal across sides: the only difference between blocks was the side where water was paired with laser pulses. As expected from signals encoding reward prediction error, suppression and activation of VTA dopamine neurons at the time of outcome had opposite effects on decisions. Suppression shifted decisions away from the side paired with laser pulses, whereas activation shifted the decisions towards this side (Figure 5E,F, *P* < 0.01, 1-way ANOVA; Figure S5H-J). Similar to the effect of manipulations of reward seen earlier, the effects of these optogenetic manipulations was strongest when the evidence was weakest (Tai et al., 2012): at low visual contrast. Dopamine-dependent psychometric shifts developed over ~8 trials after block switches (Figure 5G, Figure S5K). We observed similar psychometric shifts in experiments where we activated dopamine in a random subset of trials rather than in blocks of trials, indicating that dopamine activation in one trial is enough to influence subsequent choices (Figure S5L).

The model’s estimates of reward prediction error *δ* precisely captured the effects of these manipulations (Figure 5E-G, Figure S5H-N). To model dopamine manipulation we added to *δ* a factor that was negative for dopamine suppression and positive for dopamine activation (Figure S5M,N). This addition does not lead the model to arbitrarily low or high estimates of value, because as estimates progressively deviate from veridical, they lead to more errors, which correct the estimates towards more reasonable steady-state values (Figure S5O). The behavior of the mice conformed to these predictions: mice did not develop pure biases for one action or the other, but rather shifted their psychometric curves sideways (Figure 5E,F) and rapidly reached steady-state (Figure S5K).

## Discussion

These results reveal distinct causal roles of prelimbic cortex and of VTA dopamine neurons in learning from decisions informed by sensory confidence and reward value. We found that choices of mice reflect current sensory evidence, past rewards, and past decision confidence, and that these choices are captured by a simple reinforcement model that estimates two key internal variables. The first variable, predicted value, was causally encoded in the activity of prelimbic neurons, and non-causally reflected in the pre-outcome activity of midbrain dopamine neurons. The second variable, prediction error, was encoded in the post-outcome activity of VTA dopamine neurons. Just as in the behavioral model, both of these signals precisely depended on the confidence in the perceptual decision and on the history of reward values. Also, as in the model, these signals were necessary not for performing the ongoing trial but rather for learning from the outcome of the trial; pre-outcome activity of PL neurons and post-outcome responses of dopamine neurons drove learning.

In our task, the psychometric shifts due to past rewards are beneficial for maximizing rewards, but adjusting learning according to past decision confidence is detrimental. Indeed, stimuli were presented in random order, so they had no bearing on the next trial. Nevertheless, such confidence-dependent learning could be beneficial in natural settings where the sequence of stimuli is not random. Our experiments suggest dopamine responses as the underlying neuronal substrate for this form of learning; dopamine reward prediction errors scaled with decision confidence and psychometric shifts were most prominent when these neuronal responses were large.

Sensory confidence signals have been observed in multiple brain regions, including parietal cortex, orbitofrontal cortex, and dorsal pulvinar (Kepecs et al., 2008; Kiani and Shadlen, 2009; Komura et al., 2013), in tasks that manipulated sensory uncertainty. The signals we measured in PL during our task grew similarly with sensory confidence. Because our task also manipulated reward value, we were able to observe that PL encodes a product of this sensory confidence with reward value. This product is the predicted value of a choice. Moreover, inactivation revealed that these predicted value signals in PL play a causal role in learning: reducing them influenced future rather than ongoing choices. PL might not be the only region of frontal cortex carrying these signals. For example, similar signals necessary for learning may be observed in orbitofrontal cortex (Miller et al., 2018; Takahashi et al., 2009). Thus, the results provide evidence for the idea that manipulating predicted value, just like manipulating outcome, should result in learning driven by prediction error.

The similarities and differences that we observed in prelimbic and VTA dopamine neurons suggest how they might be functionally related. Specifically, our results suggest that dopamine neurons receive reward predictive signals from PL, and subsequently compute appropriate prediction errors. Indeed, learning was affected by pre-outcome manipulation of prelimbic neurons, but not dopamine neurons. Conversely, learning was guided by post-outcome manipulation of dopamine neurons but not prelimbic neurons. A causal role of frontal cortex in shaping dopamine responses would be consistent with anatomical projections (Beier et al., 2015; Carr and Sesack, 2000; Morales and Margolis, 2017), with simultaneous frontal-VTA recordings (Fujisawa and Buzsaki, 2011), and with recordings following pharmacological manipulations (Starkweather et al., 2018). These frontal projections may in particular affect VTA inhibitory neurons, which could then play a causal role in subtracting predicted value from observed reward (Eshel et al., 2015; Gao et al., 2007).

Neuroscience has at first studied decisions informed by perception and rewards in separate behavioral tasks, yielding data and models that are elegant yet segregated (Sugrue et al., 2005; Summerfield and Tsetsos, 2012). Our work brings these together and offers a framework for how the brain performs decisions guided by reward value and sensory evidence. This framework reveals how the brain uses both confidence and reward value for driving learning, so that learning is strongest when rewards are obtained by making a hard, low confidence, decision.

## Acknowledgements

We thank Rakesh K. Raghupathy for histology, and Kevin Miller, Nicholas Steinmetz, William Stauffer and Lauren Wool for valuable comments. This work was supported by the Wellcome Trust (grant 205093 to M.C. and K.D.H. and grant 106101 to A.L.). M.C. holds the GlaxoSmithKline/Fight for Sight Chair in Visual Neuroscience.

---

**Figure S1.**
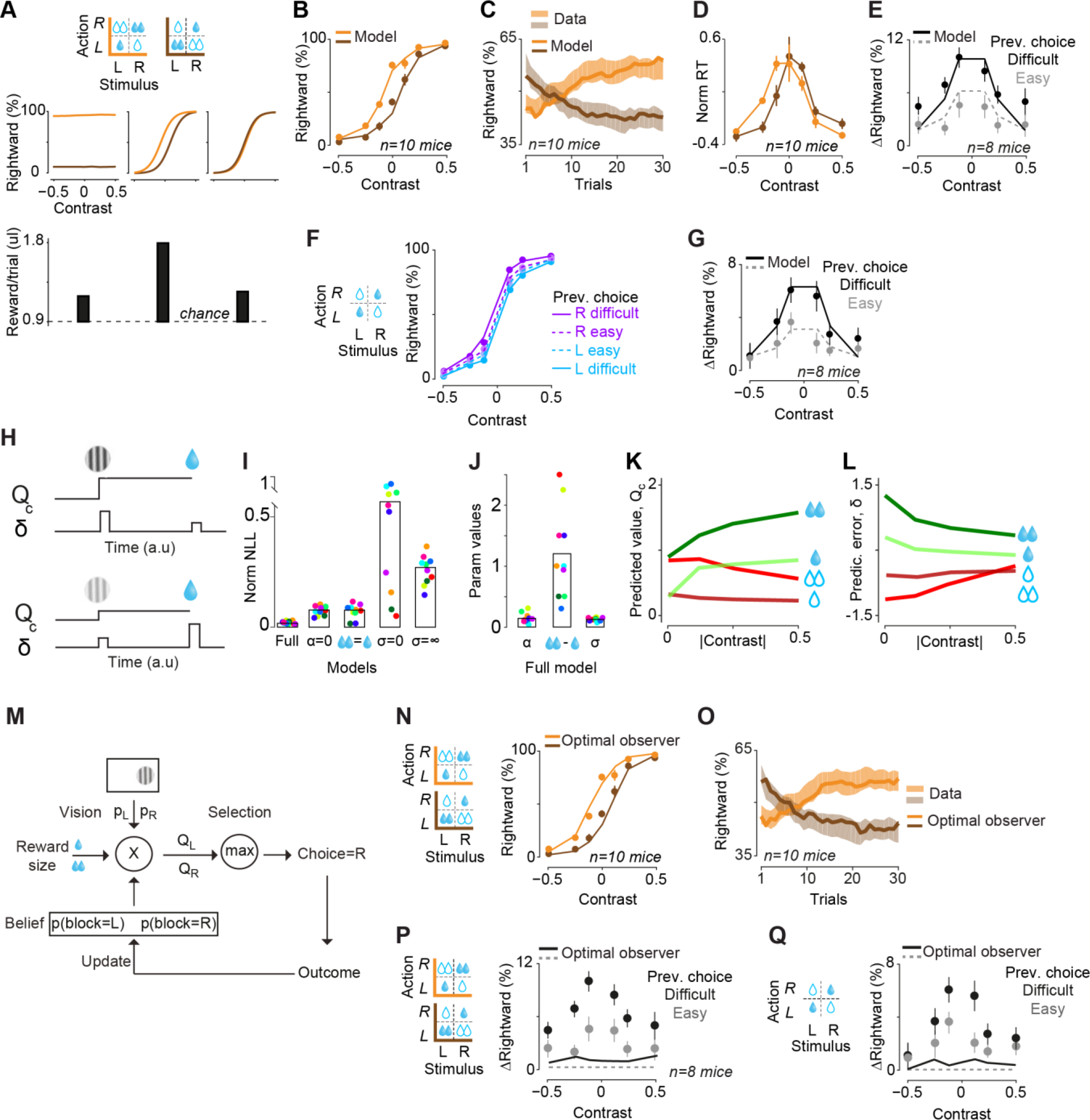
Behavioral and computational signatures of decisions guided by reward values and sensory evidence. **A**) Simulation of three agents that performed the behavioral task (top row) and their average reward harvest during 5000 trials. Left: an agent that made frequent decisions towards the side paired with larger reward, regardless of the stimulus contrast. Middle: an agent that integrated past rewards and sensory evidence. Right: an agent that made decisions only according to the sensory stimulus, ignoring the reward size. Chance level in the lower panel indicates the reward harvest of an agent that made left and right decisions in a random fashion. Simulations are performed using the model described in Figure 1. For the simulation on the left, *σ*^2^=2 and *α*=0.2 so the model could not perform the visual task but could learn from the past rewards. For the simulation in the middle panel we set *σ*^2^=0.2 and α=0.2. For the simulation on the right, we set *σ*^2^ =0.2 and *α*=0, so the model could perform visual detection but could not learn from past rewards. **B**) Average performance of mice in blocks with large reward on the left (brown) or on the right (orange). Curves are model fits on the data. **C**) Average learning curves following block switch for mice (shaded regions, mean ± s.e.) and model predictions (curves). **D**) Reaction time of animals. Reaction times were z-scored before averaging across sessions and mice. Shorter reaction times were seen for high stimulus contrast and for stimuli on the side indicating larger reward size for the current block (P < 10^−10^, 2-way ANOVA). Error bars: s.e. across mice. **E**) Average change in the proportion of rightward choices after correct difficult decisions (black) and after easy choices (gray) in the experiment with reward size manipulation. The curves are the prediction of the model. **F**) Performance of an example animal as a function of the difficulty of previous correct choices in the visual decision task with no reward manipulation. **G**) Similar to (E) but for the experiment without reward manipulation. **H**) Schematic of predicted value of choice, *Q_*C*_*, and prediction error, *δ*, of the model in trials where a high contrast or low contrast stimulus led to the same reward. **I**) Cross-validated negative log likelihood of different variants of the model. Full model contained all parameters, while each reduced model excluded one of them. Circles with different colors represent different animals. **J**) Estimated parameters of the best model, i.e. the full model. **K**) Averaged estimates of *Q*_*C*_ as a function of absolute contrast (i.e. regardless of side), for correct decisions towards the large-reward side (dark green) and correct decisions towards the small-reward side (light green), error trials towards the large-reward side (red) and error trials toward the small-reward side (dark red). **L**) Similar to (I) but for reward prediction error δ. **M**) Schematic of the optimal observer. The model uses three quantities to compute the expected value of left or right actions: the sensory evidence, the size of the rewards, and the probability (belief) that it is the left or right block. After making a choice, the model observes the outcome and updates the belief about which choice direction is associated with the larger reward. Receiving a small or large reward causes learning because they are informative about which side is paired with larger reward, but receiving no reward (error trial) is not informative (see Methods). **N**) The model accounts for the psychometric shifts in blocks with larger rewards in the left or right. **O**) The model accounts for learning curves after the block switch. **P**) The model does not account for the dependence of decisions on the difficulty of past sensory judgment. The curves are the model fits and the data are identical to those in (E). **Q**) Similar to (P) but for the task with no reward size manipulation.

**Figure S2.**
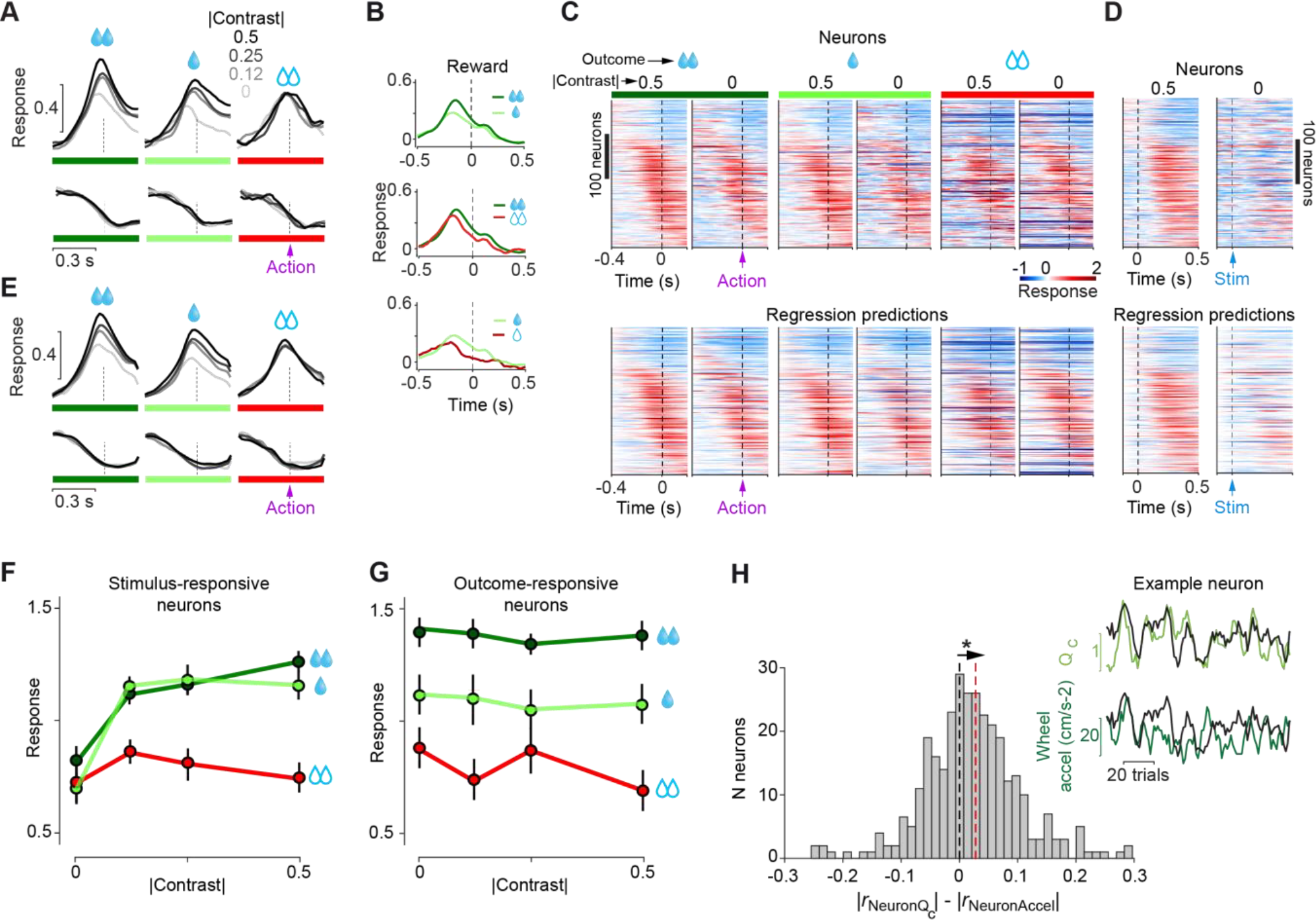
PL neuronal responses during the task. **A**) Mean population activity triggered on action onset. Responses were z-scored before averaging. Only neurons with decreased activity at the time of action were included. Left: correct choices towards the large-reward side; Middle: correct choices towards the small-reward side; Right: incorrect choices towards the large-reward side. Shades of gray: stimulus contrast. **B**) Mean population activity triggered on outcome onset. **C**) Top: Normalized responses aligned to action onset. Different panels show all cells’ responses averaged across trials of the contrast and reward value shown above the panels. Dark blue horizontal lines in error trial panels (rightmost) reflect the fact that in few recording sessions animal performed very few error trials. Bottom: Same as upper panels but for predictions of the regression that only included action events, i.e. only allowed action profiles and their coefficients. **D**) Top: normalized neuronal responses aligned to stimulus onset in all trials with |contrast| = 0.5 or 0. Bottom: these responses are accurately predicted by the regression that only included action events. **E**) Predictions of the regression triggered on action, as a function of stimulus contrast and trial type for neurons shown in (A). **F**) Average stimulus responses of neurons with significant stimulus profile as a function of stimulus contrast and reward size. Neurons which responded at the time of the stimulus encoded stimulus contrast and had lower responses in error trials than in correct trials but did not encode the size of the upcoming rewards (67/316 neurons stat), therefore did not reflect *Q*_*C*_. **G**) Average outcome responses of neurons with significant outcome profile as a function of stimulus contrast and reward size., neurons responding at the time of outcome encoded outcome value, i.e. large reward, small reward or no reward, independent of stimulus contrast (48/316 neurons, stat), and thus did not reflect δ. **H**) Correlation of PL neurons with trial-by-trial *Q*_*C*_ and trial-by-trial wheel acceleration during decision, as a proxy of response vigor. PL neurons were better correlated with *Q*_*C*_ compared to wheel acceleration (54 vs 22 neurons, P < 0.01, linear partial correlation). The inset shows the fluctuation of responses of an example neurons vs estimated *Q*_*C*_ as well as wheel acceleration during decision.

**Figure S3.**
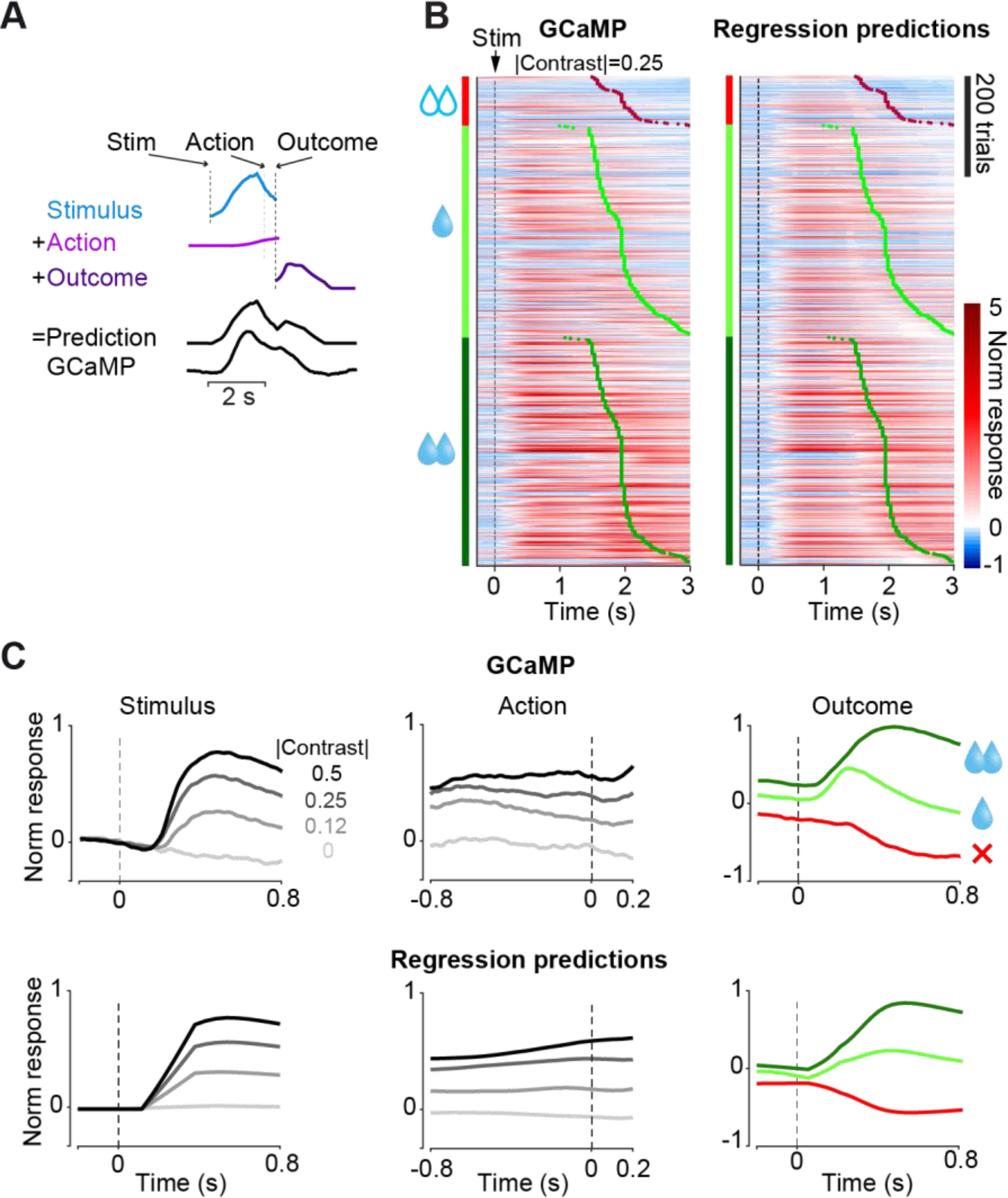
Dopamine neuronal activity during the task. **A**) Schematic of regression analysis. The regression estimates a temporal profile for each task event, which are convolved with the event times, scaled in each trial with a coefficient and summed to produce regression predictions. **B**) Left: trial-by-trial dopamine responses (as in Figure 3C), aligned to the stimulus onset (dashed line), with trials arranged vertically by trial type and outcome time (red/green points). Right: predictions of the regression analysis that only included stimulus and outcome events. **C**) Mean dopamine activity aligned to the stimulus, action and outcome. Lower panels show the predictions of the regression analysis.

**Figure S4.**
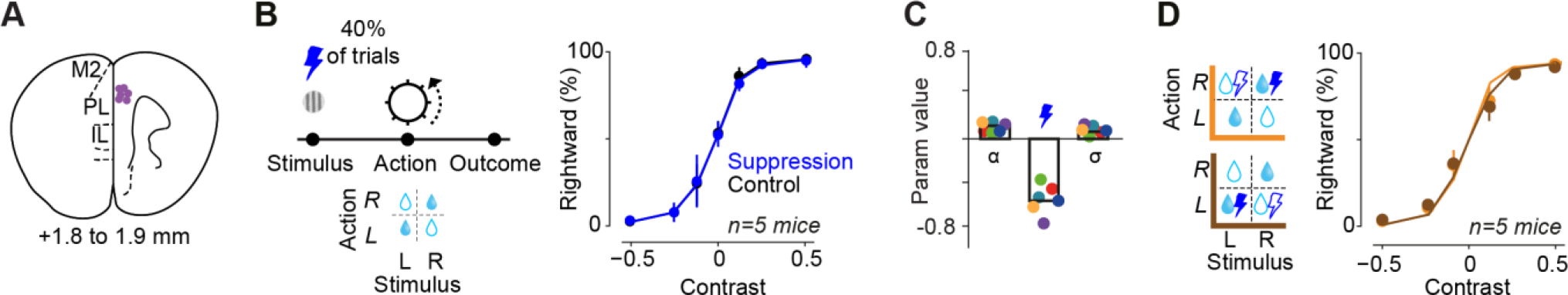
Optogenetic manipulation of prelimbic neurons during the task. **A**) Localization of implanted fiber tips from histological slices in each mouse. B) Optogenetic suppression of PL neurons at the onset of visual stimulus did not influence the psychometric curves (P = P = 0.97, 1-way ANOVA) nor their associated reaction times (P = 0.84, 1-way ANOVA). Curves are model fits. **C**) Estimated model parameters for experiments in which PL was suppressed during the stimulus presentation in the task with unequal reward size. Optogenetic suppression reduced the estimates of predicted value of choice. Circles with different colors represent different animals. **D**) Optogenetic suppression of PL neurons during the outcome time did not influence the psychometric curves (P = 0.96, 1-way ANOVA) nor their associated reaction times (P = 0.4, 1-way ANOVA).

**Figure S5.**
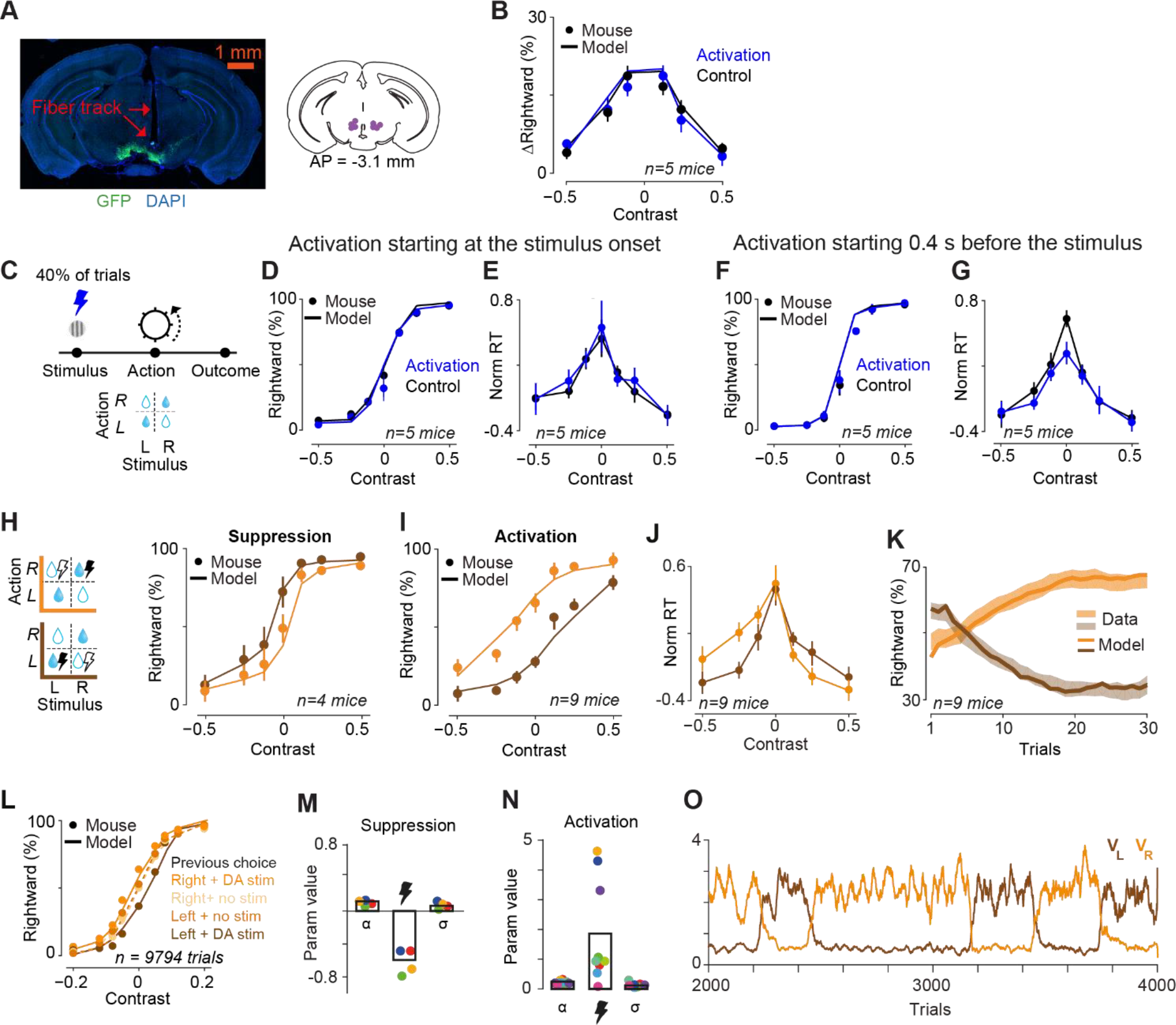
Optogenetic manipulation of VTA dopamine activity during the task. **A**) Left: Example confocal image showing optical fiber track above VTA and expression of ChR2-GFP in midbrain of DAT-Cre mouse. Right: localization of implanted fiber tips from histological slices in all mice. **B**) Activation of dopamine neurons during stimulus presentation in the task with reward manipulation. Activation did not change the shifts of psychometric curves. **C**) Activation of dopamine neurons during stimulus presentation in a task with no reward manipulation. Optogenetic activations were applied in 40% of randomly chosen trials. The laser pulses started either at the stimulus onset (D,E) or 0.4 s before the stimulus (F,G). **D**) Activating dopamine neurons at the time of stimulus onset did not influence current choices (P = 0.96, 1-way ANOVA) or next trials (P = 0.9, 1-way ANOVA). **E**) Activating dopamine neurons at the time of stimulus onset did not influence reaction times (P = 0.67, 1-way ANOVA). **F**) Activation of dopamine neurons 0.4 s prior to stimulus presentation did not affect ongoing decisions (P = 0.99, 1-way ANOVA) or subsequent decisions (P = 0.80, 1-way ANOVA). **G**) Activating dopamine neurons prior to stimulus onset mildly decreased reaction times, which barely reached statistical significance in in trials with contrast=0 (P=0.05, singed rank test). **H**) Left: Suppression or activation of dopamine during the reward time. In consecutive blocks of trials, correct choices to left or right were paired with the laser pulses. Right: The effect of optogenetic suppression. **I**) The effect of optogenetic activation. **J**) Reaction times separated for stimuli and dopamine activation blocks. Reaction times were smallest for high-contrast stimuli and for response side paired with dopamine. **K**) Mean learning curves following switch of dopamine activation side. Shaded regions: mean ± s.e.; curves: model prediction. **L**) Performance of an example mouse in experiments in which dopamine neurons were activated in randomly chosen successful trials (~30% of trials). Psychometric functions are plotted as a function of the side chosen on the previous trial, and outcome of that trial. Activation of dopamine neurons in previous trial shifted decisions towards the side paired with such activation, but only when the immediate sensory evidence was weak. Solid curves: predictions of the behavioral model. **M**) Estimated model parameters for experiments in which dopamine neurons were suppressed in consecutive blocks of trials. Circles with different colors represent different animals. **N**) Similar to (E) but for the experiments including activation of dopamine neurons in consecutive blocks. **O**) Simulation showing *V*_*L*_ and *V*_*R*_ over trials in a model run in which we added a constant (=3) to the value of δ in correct trials of alternating blocks. *V*_*L*_ and *V*_*R*_ stay stable since rewards are contingent to correct sensory detection.

## Methods

All experiments were conducted according to the UK Animals Scientific Procedures Act (1986) under appropriate project and personal licences.

### Animals and surgeries

The data presented here was collected from 33 mice (19 male) aged between 10-24 weeks. All mice were first implanted with a custom metal head plate. To do so, the animals were anesthetized with isoflurane, and were kept on a feedback-controlled heating pad (ATC2000, World Precision Instruments, Inc.). Hair overlying the skull was shaved and the skin and the muscles over the central part of the skull were removed. The skull was thoroughly washed with saline, followed by cleaning with sterile cortex buffer. The head plate was attached to the bone posterior to bregma using dental cement (Super-Bond C&B; Sun Medical). For electrophysiological experiments, we covered the exposed bone with Kwik-Cast (World Precision Instruments, Inc.), trained the animals in the behavioral task in the following weeks, and subsequently performed a craniotomy over the frontal cortex for lowering the silicon probes. For fiber photometry and optogenetic experiments, after the head plate fixation, we made a craniotomy over the target area (PL or VTA) and injected viral constructs followed by implantation of the optical fiber, which was secured to the head plate and skull using dental cement. Post-operative pain was prevented with (Rimadyl) on the three following days.

### Behavioral tasks

Behavioral training started at least 7 days after the head plate implantation surgery. For mice which received viral injection, training started 2 weeks after the surgery. Animals were handled and acclimatized to head fixation for 3 days, and were then trained in a 2-alternative forced choice visual detection task (Burgess et al., 2017). After the mouse kept the wheel still for at least 0.5 s, a sinusoidal grating stimulus of varying contrast appeared on either the left or right monitor, together with a brief tone (0.1 s, 12 kHz) indicating that the trial had started. The mouse could immediately report its decision by turning the wheel located underneath its forepaws. Wheel movements drove the stimulus on the monitor, and a reward was delivered if the stimulus reached the center of the middle monitor (a correct trial), but a 2s white noise was played if the stimulus reached the center of the either left or right monitors (an error trial). The inter trial interval was set to 3 s. As previously reported, well-trained mice often reported their decisions using fast stereotypical wheel movements (Burgess et al., 2017). In the initial days of the training (first 4 to 7 days), stimuli had contrast=1. Lower-contrast stimuli were introduced when the animal reached the performance of ~ 70%. After 2-3 weeks of training, the task typically included 7 levels of contrast (3 on the left, 3 on the right and zero contrast) which were presented in a random order across trials with equal probability. We finally introduced unequal water rewards for correct decisions: in consecutive blocks of 50-350 trials (drawn from a uniform distribution), correct decisions to one side (left or right) were rewarded with larger reward (2.4 μl vs 1.2 μl of water) (Figure 1).

Experiments involving optogenetic manipulation of PL neurons or VTA dopamine neurons had the same timeline as described above (Figure 4,5). In experiments involving fiber photometry, the task timeline slightly differed from above, allowing longer temporal separation of stimulus, action and outcome (Figure 3). In these experiments, wheel movements immediately after the visual stimulus did not move the stimulus on the monitor and did not result in a decision (open-loop condition). Instead, an auditory go cue (0.1s) which was played 0.6-1.8 s after the stimulus onset started the closed-loop during which animals could report the decision. Wheel movements prior to go cue did not terminate the trial and we did not exclude these trials from our analysis (excluding these trials did not affect our results). In these experiments, we defined the action time as the onset of first wheel movement after the stimulus onset. In all experiments, reaction times were measured from the onset of visual stimulus till the onset of the first wheel movement.

The behavioral experiments were controlled by custom-made software written in Matlab (Mathworks) which is freely available at: github.com/cortex-lab/signals. Instructions for both the software as well as hardware assembly is freely accessible at: www.ucl.ac.uk/cortexlab/tools/wheel.

### Electrophysiological experiments

We recorded neuronal activity in prelimbic frontal cortex (PL) using multi-shank silicon probes in wild-type C57/BL6J mice. We implanted the animals after they fully learned to perform the task, performing the final stage of the behavioral task (including block switches) with performance above 70% for at least three sessions. A 32-channel, 2 shank silicon probe (Cambridge NeuroTech) was mounted on a moveable miniature Microdrive (Cambridge NeuroTech) and implanted it into PL (n=6 mice). On the implantation day, we removed the Kwik-Cast cover from the skull and drilled a small incision in the cranium over the frontal cortex, ML = 0.3 mm, AP = 1.8 mm (burr #19007–07, Fine Science Tools). The brain was protected with Ringer solution. We lowered the probe through the intact dura using a manipulator (PatchStar, Scientifica) to 1.4 mm from the dura surface. The final approach towards the target depth (the last 100–200 μm) was performed at a low speed (2–4 μm/sec), to minimize potential damage to brain tissue. Once the probe was in its required position, we waited 10 minutes to let the brain recover from the insertion and fixed the Microdrive on the head plate using dental cement. For reference signal we used a skull screw implanted on the skull ~ 3-4 mm posterior to the recording site. At the end of each recording day we lowered the Microdrive 100 μm.

Recordings were performed using OpenEphys system. Broadband activity was sampled at 30 kHz (band pass filtered between 1 Hz and 7.5 kHz by the amplifier) and stored for offline analysis. Recorded spikes were sorted using the KlustaSuite software (Rossant et al., 2016) (www.cortexlab.net/tools), which involves three packages, one for each of three steps: spike detection and extraction, automatic spike clustering, and manual verification and refining of the clusters. Manual spike sorting was performed oblivious to task-related responses of the units.

### Fiber photometry experiments

To measure the activity of dopamine neurons, we employed fiber photometry (Gunaydin et al., 2014; Lerner et al., 2015). We injected 0.5 µL of diluted viral construct (AAV1.Syn.Flex.GCaMP6m.WPRE.SV40) into the VTA:SNc (ML:0.5 mm from midline, AP: −3 mm from bregma and DV:-4.4 mm from the dura) of DAT-Cre mice backcrossed with C57/BL6J mice (B6.SJLSlc6a3tm1.1(cre)Bkmn/J). We implanted an optical fiber (400 µm, Doric Lenses Inc.) over the VTA, with the tip 0.05 mm above the injection site. We used a single chronically implanted optical fiber to deliver excitation light and collect emitted fluorescence. We used multiple excitation wavelengths (465 and 405 nm) modulated at distinct carrier frequencies (214 and 530 Hz) to allow for ratiometric measurements. Light collection, filtering, and demodulation were performed as previously described (Lerner et al., 2015) using Doric photometry setup and Doric Neuroscience Studio Software (Doric Lenses Inc.). For each behavioral session, least-squares linear fit was applied to the 405nm control signal, and the ΔF/F time series was then calculated as ((490nm signal – fitted 405nm signal) / fitted 405nm signal). All analyses were done by calculating *z*-scored ΔF/F.

### Optogenetic experiments

#### Optogenetic manipulation of PL neurons

For suppressing PL responses, we injected 0.5 µL of diluted viral construct containing ChR2 (AAV5.EF1a.DIO.hChr2(H134R)-eYFP.WPRE) unilaterally into the PL (ML:0.3 mm, AP: 1.8 mm from bregma and DV:-1.6 mm from the dura) of Pvalb-Cre mice backcrossed with C57/BL6J (B6.129P2-Pvalb^tm1(cre)Arbr^/J). We implanted an optical fiber (200 µm, Doric Lenses Inc.) over the PL, with its tip staying 0.4 mm above the injection site. We waited 2 weeks for virus expression and then started the behavioral training. After achieving stable task performance using symmetric water rewards, we introduced laser pulses which had following parameters: 473 nm (Laserglow LTD), number of pulses: 12, each pulse lasting 10 ms and separated by 30 ms, laser power: ~2-3 mW (measured at the fiber tip). The laser pulses were applied either from the stimulus onset (Figure 4, Figure S4) or during the outcome (Figure S4). Manipulation at the time of the stimulus included three types of experiments: a) in 40% of randomly chosen trials in the task that had blocks of 50-350 trials with unequal rewards, b) in the task that had blocks of 50-350 trials with unequal rewards each of them with or without laser pulse at the stimulus time, making four types of blocks, c) in 40% of randomly chosen trials of a purely visual task (with symmetric and stable rewards). In the experiments involving manipulations at the trial outcome, in consecutive blocks of 50-350 trials, correct decisions to one side, L or R, were paired with laser pulses (Figure S4).

#### Optogenetic manipulation of VTA dopamine neurons

For activating or suppressing dopamine neurons, We injected 0.5 µL of diluted viral constructs containing ChR2 (AAV5.EF1a.DIO.hChr2(H134R)-eYFP.WPRE) or Arch3 (rAAV5/EF1a-DIO-eArch3.0-eYFP) unilaterally into VTA:SNc (ML:0.5 mm from midline, AP: −3 mm from bregma and DV:-4.4 mm from the dura) of DAT-Cre mice backcrossed with C57/BL6J mice (B6.SJLSlc6a3tm1.1(cre)Bkmn/J). We implanted an optical fiber (200 µm, Doric Lenses Inc.) over the VTA, with its tip staying 0.4 mm above the injection site. We waited 2 weeks for virus expression and then started the behavioral training. After achieving stable task performance using symmetric water rewards, we introduced laser pulses which had the following parameters: 473 nm and 532 nm for ChR2 and Arch3, respectively (Laserglow LTD), number of pulses: 12, each pulse lasting 10 ms and separated by 30 ms, laser power: ~8 mW (measured at the fiber tip). For the suppression experiment using Arch3, in few sessions we used a single 300 ms long pulse. The laser pulses were applied either 0.4 s prior to the stimulus (Figure S5), exactly at the time of the stimulus (Figure 5, Figure S5), or at the time of the reward (Figure 5, Figure S5). For experiments involving activation of dopamine neurons prior to the stimulus onset, in 40% of randomly chosen trials, we delivered laser pulses. For experiments involving activation of dopamine neurons at the stimulus onset, we either applied pulses in 40% of randomly chosen trials (Figure S5) or in blocks of 50-350 trials (Figure 5). In the experiments involving manipulation of dopamine activity at the trial outcome, in consecutive blocks of 50-350 trials, correct decisions to one side, L or R, were paired with laser pulses (Figure 5). In experiments involving trial-by-trial manipulations at the trial outcome (rather than blocks of trials), in 30% of randomly chosen correct trials, the reward was paired with laser pulses (Figure S5). In both these experiments the laser was turned on simultaneously with the TTL signal that opened the water valve.

### Histology and anatomical verifications

To verify expression of viral constructs we performed histological examination. Animals were deeply anesthetized and perfused, brains were post-fixed, and 60 µm coronal sections were collected. For optogenetic experiments on PL, we immunostained with antibody to eYFP and secondary antibodies labelled with Alexa Fluor 488 (Figure 4). For experiments on dopamine neurons (both photometry and optogenetic), sections were immunostained with antibody to TH and secondary antibodies labelled with Alexa Fluor 594. For animals injected with ChR2 or Arch3 constructs into the VTA, we also immunostained with an antibody to eYFP and secondary antibodies labelled with Alexa Fluor 488 (Figure 5, Figure S5). We confirmed viral expression in all animals with ChR2 injections into the PL and in 14 (out of 15) mice injected with ChR2, Arch3 or GCaMP6M.

The anatomical location of implanted optical fibers was determined from the tip of the longest fiber track found, and matched with the corresponding Paxinos atlas slide (Figure 3-5, Figure S4 and S5). To determine the position of silicon probes in PL, coronal sections were stained for GFAP and matched to the corresponding Paxinos atlas (Figure 2A). Confocal images from the sections were obtained using Zeiss 880 Airyscan microscope.

### Behavioral modeling

To estimate the hidden variables that could underlie learning and decisions in our tasks, we adopted a reinforcement learning model which we developed previously (Lak et al., 2017). In our task, knowing the state of the trial (L or R) is only partially observable, and it depends on the stimulus contrast.

In keeping with the standard psychophysical treatments of sensory noise, the model assumes that the internal estimate of the stimulus, 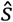, is normally distributed with constant variance around the true stimulus contrast: 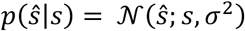. In the Bayesian view, the observer’s belief about the stimulus *s* is not limited to a single estimated value 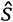. Instead, 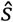 parameterizes a belief distribution over all possible values of *s* that are consistent with the sensory evidence. The optimal form for this belief distribution is given by Bayes rule:

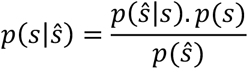

We assume that the prior belief about *s* is uniform, which implies that this optimal belief will also be Gaussian, with the same variance as the sensory noise distribution, and mean given by 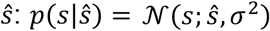. From this, the agent computes a belief, i.e. the probability that the stimulus was indeed on the right side of the monitor, 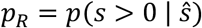, according to:

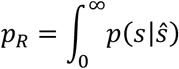

*p*_*R*_ represents the trial-by-trial probability of the stimulus being on the right side (and *p*_*L*_ = 1 − *p*_*R*_ represents the probability of it being on the left).

The expected values of the two choices L and R are computed as *Q*_*L*_ = *p*_*L*_ *V*_*L*_ and *Q*_*R*_ = *p*_*R*_*V*_*R*_, where *V*_*L*_ and *V*_*R*_ represent the stored values of L and R actions. To choose between the two options, we used an argmax rule which selects the action with higher expected value deterministically (Figure 1). Using other decision functions such as softmax did not substantially change our results. The outcome of this is thus the choice (L or R), its associated confidence *p*_*C*_, and its predicted value *Q*_*C*_.

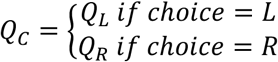

When the trial begins, i.e. when the auditory cue indicates that the trial has started, the expected reward prior to any information about the stimulus is *V*_*onset tone*_ = (*V*_*L*_ + *V*_*R*_)/2. Upon observing the stimulus and making a choice, the prediction error signal is: *Q*_*C*_ − *V*_*onset tone*_. After receiving the reward, *r*, the reward prediction error is *δ* = *r* − *Q*_*C*_.

Given this prediction error the value of the chosen action will be updated according to:

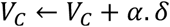

where *α* is the learning rate. For simplicity, the model does not include temporal discounting.

The model’s estimates of both *Q*_*C*_ and *δ*, depend on stimulus contrast, reward size, and whether the choice is correct (Figure 1, Figure S1). *Q*_*C*_ grows with the stimulus contrast as well as the size of reward. Perhaps less intuitively, however, the dependence of *Q*_*C*_ on contrast is reversed on error trials (Figure 1J, *red curve*). This effect is easily understood if *V*_*L*_ = *V*_*R*_. In this case, errors are entirely due to wrong sensory estimates of *p*_*L*_ and *p*_*R*_. If a stimulus is on the R, the observer chooses L only if *p*_*c*_ = *p*_*L*_ > *p*_*R*_. In high-contrast trials, this occurs rarely and by a small margin (Kepecs and Mainen, 2012; Lak et al., 2017), so *p*_*c*_ ≈ 0.5 and *Q*_*C*_ is low. At lower contrast, instead, this can occur more often and with *p*_*c*_ ≫ 0.5, so *Q*_*C*_ is higher.

### Model fitting

The experiments included sessions with blocks of trials with unequal water rewards and sessions with no reward size manipulation. In the optogenetic experiments, these sessions could include suppression of PL neurons or activation/suppression of VTA dopamine neurons.

We fitted our model as well as reduced model variants on choices acquired in the task with unequal water rewards and cross-validated the necessity of model parameters. We then used the model that could best account for the data and fitted it on the experiments that included optogenetic manipulations.

#### Experiments with unequal water reward

For fitting, we set the value of smaller water reward to 1. Thus, the payoff matrix for blocks with larger reward on the left or right, respectively, are:

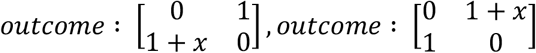

where *x*, a constant, represents the value of extra drop of water. We set the payoff for incorrect decisions to zero in all our model fitting.

We fitted the model as well as reduced model variants on the decisions of mice in the task with unequal water rewards, and cross-validated the necessity of model parameters (Figure S1). As described above, the full model included the following parameters: σ^2^, *x*, *α*. Each reduced model did not include one of these parameters. For σ^2^, one reduced model was set to have σ^2^ = 0, representing a model with no sensory noise, and the other reduced model was set to have σ^2^ = ∞, representing a model with extremely large sensory noise. For cross-validated fitting, we divided sessions of each mouse to 3 and performed a 3-fold cross validation. We performed the fit and parameter estimation on the training sessions and used the estimated parameters against the test sessions for computing goodness of fit. For fitting, we performed exhaustive search in the parameter space expanding large value range for each of the parameters to find the best set of model parameters that account for the observed decisions. We searched the following parameter space: *α* = 0: 0.05: 0.95, σ^2^ = 0.04: 0.04: 0.8 and *x* = −5: 0.2: 10. To do so, for each possible combination of these parameters, we repeatedly fed the sequences of stimuli that each mouse experienced to the model, observed decisions (iteration = 1000), and averaged across the iterations to compute the probability that model made a leftward and rightward decision 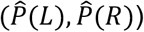 for each trial. We then calculated the negative log likelihood (NLL) as the average of 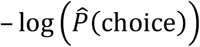, where choice indicates the mouse’s decision in each trial (Figure S1). The set of parameters that gave the lowest NLL were used to compute goodness of fit in the test sessions (3-fold cross-validation).

#### Manipulation of PL activity

For experiments including suppression of the PL at the stimulus onset, we allowed the model to add a constant to the predicted value of the choice *Q*_*C*_. A negative constant resulted in lower predicted value and hence increased prediction error after receiving a reward (Figure S4). We fitted the model on choices as described above.

#### Manipulation of dopamine activity

For experiments including suppression or activation of dopamine neurons at the outcome time, we allowed the model to add a constant to the reward prediction error *δ*. This constant was negative for the experiment with dopamine suppression, and was positive for the experiment with dopamine activation (Figure S5). We fitted the model on choices as described above.

#### Optimal observer model fitting

We constructed an alternative class of model that optimally performs our task. This observer leverages the structure of the task, i.e. it knows that only two reward sizes are available and that they switch side occasionally. The observer would thus only need to infer whether it is in the left or the right block, given the sequence of outcomes in the previous trials. To do so we used a hidden Markov model (Matlab HMM toolbox). The model estimates the trial-by-trial probability that the current state is left or right block *p*_(*S*=*L*)_ and *p*_(*S*=*R*)_, respectively, given a state transition matrix and an observation matrix. The state transition matrix defines the probability of block switch, which can be calculated from the number of block switches and number of trials in each dataset. The observtion matrix defines the probability of observed outcomes (no reward, small reward and larger reward) given each state. The model computes the expected value of left and right actions according to:

*Q*_*L*_ = *p*_*L*_(*p*_(*S*=*L*)_*r*_(*a*=*L*,*s*=*L*)_ + *p*_(*S*=*R*)_*r*_(*a*=*L*,*s*=*R*))_ and *Q*_*R*_ = *p*_*R*_(*p*_(*S*=*L*)_*r*_(*a*=*R*,*s*=*L*)_ + *p*_(*S*=*R*)_*r*_(*a*=*R*,*s*=*R*)_), where *p*_*L*_and *p*_*R*_ are estimated as described in the reinforcement learning model section, *r*_(*a*=*L*,*s*=*L*)_ indicates the size of reward available for left action in the L block and *p*_(*S*=*L*)_ and *p*_(*S*=*R*)_ are the probabilities that the current trial belongs to L or R block, estimated using the hidden Markov model. This model learns about the blocks from any reward trials (both small and large rewards). This learning is, however, not influenced by the sensory confidence. When emission matrix set optimally (i.e. in the left block the probability of large reward on the right to be zero, *p*(*r* = *large*|*a* = *R*, *s* = *L*) = 0, etc.), the model learns the block switch after only one rewarded trial. However, we observed that mice took several trials to learn the block switch (Figure S1). Thus, for the fitting purpose, we considered that the observation matrix is noisy (*p*(*r* = *large*|*a* = *R*, *s* = *L*) = *β*). An intuition behind this could be that the mouse does not always accurately detect the size of reward, and is hence slightly confused about the size of rewards which are available for L and R choices in each block; *β* determines this noise level. We estimated *β* for each animal using exhaustive search, as described in the previous section. After fitting, the model could account for the dependence of decisions on past rewards and current sensation (Figure S1), but not for the dependence of choices on decision confidence in the previous trial (Figure S1).

### Neuronal regression analysis

In order to quantify how each task event (stimulus, action, outcome) contributes to neuronal activity, and, the extent to which trial-by-trial variation in neuronal responses reflect animal’s estimate of pending reward and prediction error, we set up a neuronal response model (Park et al., 2014) (Figure 2,3 S2, S3).

We modelled the spiking activity of a neuron during trial *j*, which we denote *R*_*j*_(*t*) as

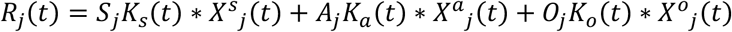

In the above equation, *K*_*s*_(*t*), *K*_*a*_(*t*) and *K*_*o*_(*t*) are the profiles (kernels) representing the response to the visual stimulus, the action, and the outcome. *X*^*s*^_j_ (*t*), *X*^*a*^_j_ (*t*) and *X*^*o*^_j_ (*t*) are indicator functions which signify the time point at which the stimulus, action and outcome occurred during trial *j*. *S*_*j*_, *A*_*j*_ and *O*_*j*_ are multiplicative coefficients which scale the corresponding profile on each trial and * represents convolution. Therefore, the model represents neuronal responses as the sum of the convolution of each task event with a profile corresponding to that event, which its size was scaled in each trial with a coefficient to optimally fit the observed response. Given the temporal variability of task events in different trials, the profile for a particular task event reflects isolated average neuronal response to that event with minimal influence from nearby events. The coefficients provide trial-by-trial estimates of neuronal activity for each neuron.

The model was fit and cross-validated using an iterative procedure, where each iteration consisted of two steps. In the first step the coefficients *S*_*j*_, *A*_*j*_ and *O*_*j*_ were kept fixed and the profile shapes were fitted using linear regression. Profiles were fitted on 80% of trials and were then tested against the remaining 20% test trials (5-fold cross-validation). In the second step, the profiles were fixed and the coefficients that optimized the fit to experimental data were calculated, also using linear regression. Five iterations were performed. In the first iteration, the coefficients were initialized with value of 1. We applied the same analysis on the GCaMP responses (Figure 3, Figure S3).

We defined the duration of each profile to capture the neuronal responses prior to or after that event and selected longer profile durations for the GCaMP data to account for Ca^+2^ transients (PL spike data: stimulus profile: 0 to 0.6 s, action profile: −0.4 to 0.2 s, outcome profile: 0 to 0.6 s; GCaMP data: stimulus profile: 0 to 2 s, action profile: −1 to 0.2 s, outcome profile: 0 to 3s, where in all cases 0 was the onset of the event). For both spiking and GCaMP data, the neuronal responses were averaged using a temporal window of 20 and 50 ms, respectively, and were then z-scored.

